# StoatyDive: Evaluation and Classification of Peak Profiles for Sequencing Data

**DOI:** 10.1101/799114

**Authors:** Florian Heyl, Rolf Backofen

## Abstract

The prediction of binding sites (peak calling) is a common task in the data analysis of methods such as crosslinking or chromatin immunoprecipitation in combination with high-throughput sequencing (CLIP-Seq, ChIP-Seq). The predicted binding sites are often further analyzed to predict sequence motifs or structure patterns as an example. However, the obtained peak set can vary in their profile shapes because of the used peakcaller method, different binding domains of the protein, protocol biases, or other factors. Thus, a tool is missing that evaluates and classifies the predicted peaks based on their shapes. We hereby present StoatyDive, a tool that can be used to filter for specific peak profile shapes of sequencing data such as CLIP and ChIP. StoatyDive therefore fine tunes downstream analysis steps such as structure or sequence motif predictions and acts as a quality control.

With StoatyDive we were able to classify distinct peak profile shapes from CLIP-seq data of the histone stem-loop-binding protein (SLBP). We show the potential of StoatyDive, as a quality control tool and as a filter to pick different shapes based on biological or methodical questions.

StoatyDive is open source and freely available under GLP-3 at https://github.com/BackofenLab/StoatyDive and at bioconda https://anaconda.org/bioconda/stoatydive.

## Introduction

The biological function of a protein is determined by its interaction partners and the mode of interaction. Studying these interactions broadens our horizon about the cellular mechanisms such as alternative splicing and post-transcriptional regulation. Crosslinking, or chromatin immunoprecipitation in combination with high-throughput sequencing (CLIP-Seq, ChIP-Seq) are methods to fathom these interactions. CLIP-Seq investigates all interactions between an RNA binding protein (RBP) and its target RNAs (1). CLIP-Seq thus scrutinizes the post-transcriptional regulation by RBPs. Prediction of binding regions (peak calling) is a crucial step in the data analysis of methods such as CLIP-Seq, or ChIP-Seq. Typically before the peak analysis there is no evaluation and classification of the peak characteristics. Yet, the obtained peak set from a peakcaller might have different peak profiles that are worth to filter to refine a downstream analysis. The different peak shapes are the result of several biological and technical problems.

Jankowsky and Harris (2) have discussed the characteristics of RNA-protein interactions and the potential problems. An RBP can have different bindings domains, or be part of a protein complex. Thus, the protein might have disparate binding sites with different affinities (mechanisms) ranging from specific to unspecific. Factors such as the affinity of the protein for the RNA site, the concentration of the protein and RNA influence the binding specificity of the different protein binding domains. For example, they describe mRNA export factors that have the ability to bind several RNAs. Something not mentioned is the possibility that the protein type might also manifest in different peak profiles. A helicase might have different peak profiles compared to a transcription factor.

In addition, technical biases might change the peak profile landscape. Errors during the read library preparation might introduce unspecific bindings. Protocol biases, for example, PAR-CLIP biases that are introduced by endonuclease and photoactivatable nucleoside (3), might also affect the binding site predictions. On top, the peakcaller itself might generate specific peak profiles and false positives, which the user might not want to have in their data.

That is why, many questions occur in the data analysis of binding sites. Does my protein of interest bind generally more specific (Figure 1a) or more unspecific (Figure 1b)? Does my RBP of interest have more than one binding site? Does my experiment have some quality issues, meaning, does my reads come from unspecific bindings because of library preparation errors? Does my protocol generate biases? Do I have false positives in the set of predicted peaks from my peakcaller of choice? An initial clustering approach was done by Cremona et al. (4), but only for ChIP-Seq data and only on predefined features.

**Fig. 1.**
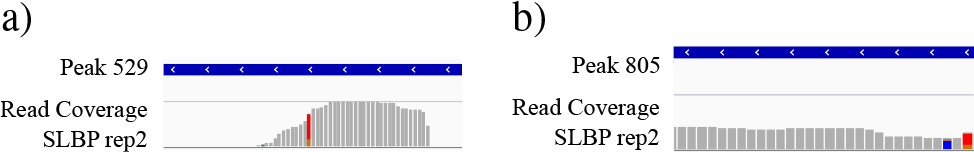
Often the peaks from a peakcaller are further inspected and evaluated based on their biological and technical reasoning. Searching for more (a) specific and more (b) unspecific binding events (peak profiles) is often done on a handful of selected peaks, because a general tool is missing. Thus, evaluating and classifying peaks is an open problem in binding site prediction to filter and thus refine down-stream analysis tasks for data such as CLIP or ChIP. StoatyDive assists to find and distinguish peaks like (a) and (b).

We hereby present StoatyDive, a tool to evaluate and classify peak profiles to help to answer the aforementioned questions. StoatyDive uses the whole peak profiles as well as predefined features to do a peak shape clustering for sequencing data. In this paper, we will test StoatyDive on CLIP data of the eCLIP protocol from the histone stem-loop-binding protein (SLBP) (5). SLBP has been reported to be a histone mRNA export and translation factor (6). StoatyDive delivers several plots and a table to assess the different binding profiles of a protein. The tool assists to select specific and unspecific binding sites and to find similar shaped peak profiles. Thus, we try to refine the obtained peaks of the SLBP data to find more specific sites of SLBP. It also helps as a quality assessment to validate a CLIP-Seq, or any other binding experiment. StoatyDive (https://github.com/BackofenLab/StoatyDive; https://anaconda.org/bioconda/stoatydive) comes with some test data and a quick installation guide.

## Algorithm

### Peak Correction, Extension and Coverage Calculation

StoatyDive was implemented in python (>= 3.6) and R (>= 3.4.4). The tool needs three files: the predicted binding regions of a peak calling algorithm in bed6 format, a bam or bed file that was used for the peak calling (experiment or control), and a tabular file of the chromosome size of the reference genome (Figure 2).

**Fig. 2.**
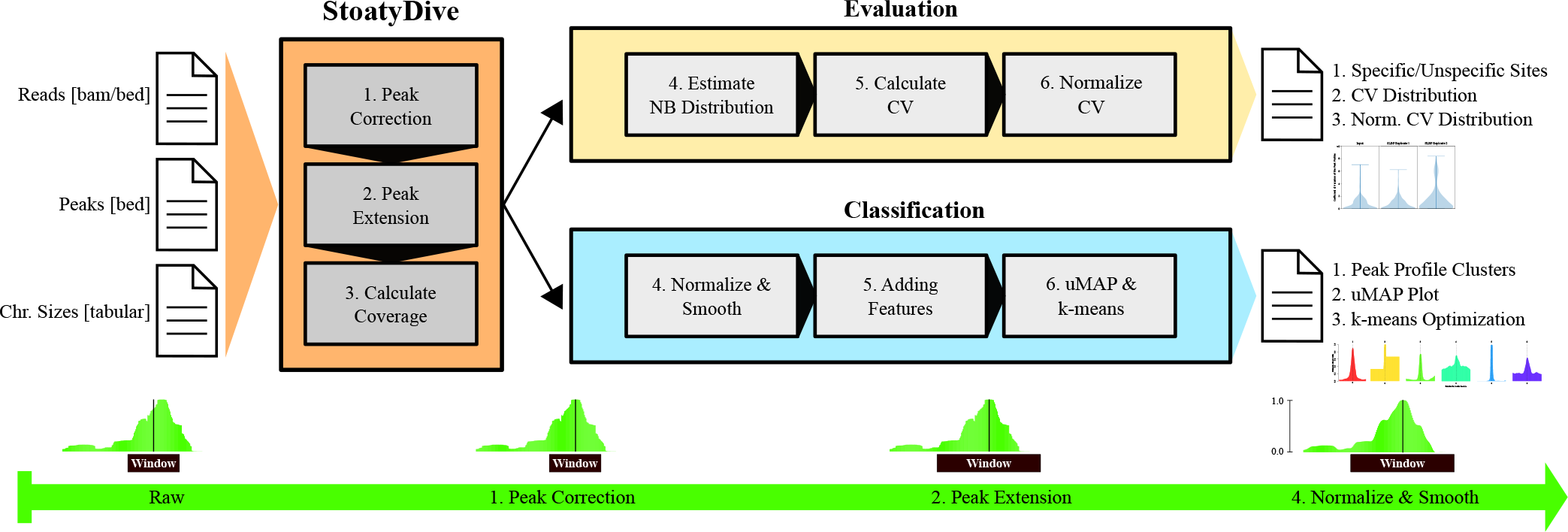
StoatyDive assists to find peaks like figure 1a or figure 1b. It has two modules: the evaluation and the classification of peak profiles. The user has to provide reads (or events), peaks and a chromosome size file. StoatyDive then shifts the peaks to their correct center (peak correction), extents the peaks to a common length (maximal peak length of peak set or user defined value), and calculates the coverage with bedtools (7). The peak correction can be turned off. In the evaluation, StoatyDive then estimates the read coverage as a negative binomial. From the hyperparameters it calculates the coefficient of variation (CV) and normalizes it (Equations 1 and 2). The normalized CV can then be used to divide the peaks into specific and unspecific sites. Furthermore, the CV distribution acts as a quality control between control and signal experiments. In the classification, StoatyDive first normalizes the peak profiles to remove the intensity as a feature. Then it smooths the profiles to support the data assumptions of uMAP (8) and to remove some noise. After that, it adds curve specific features to the data. The higher dimension of the data is then reduced with uMAP. StoatyDive then clusters the new data with k-means (9). The user then obtains several plots and a table to investigate the different peak profile clusters.

First, StoatyDive checks if a peak profile needs to be centered (peak correction). In the default mode, the profiles are centered by a convolution with a standard normal distribution. The maximum value of the convolution gives the nucleotide shift of the peak profile to center the peaks. So the window with the peak length is shifted to the center of the peak (Figure 2 step 1). With this approach we retain the context and take care of two problems. Problem one, peakcallers often produce peaks that are not correctly centered. Problem two, dimensional reduction methods, such as uniform manifold approximation and projection for dimension reduction (uMAP) (8), are not translation invariant. Thus, two profiles with the same shape but in different position might end up in different locations in the new dimensional space.

After the peak correction, StoatyDive extents the peaks by default to the maximal peak length of the given peak set (Figure 2 step 2). This removes the peak length as a potential feature for the evaluation and classification. StoatyDive then calculates the read coverage (Figure 2 step 3) for each position inside a peak with the help of bedtools (7).

### Evaluation of Peak Profile

With the results of bedtools, StoatyDive evaluates every peak *i* from the total set of *k* peaks. StoatyDive will estimate the read count for every peak as a negative binomial *X*_*i*_ ~ NB(*r*_*i*_, *p*_*i*_) with the hyperparameters *r*_*i*_ (number of hits) and *p*_*i*_ (probability of a hit). It then calculates the coefficient of variation (CV) for every peak. A simple estimation of the variance is not enough because the profile depends on the read coverage. Thus, to be able to compare each peak profile we have to normalize for the expected number of reads to adjust the variance. So the CV for each peak,

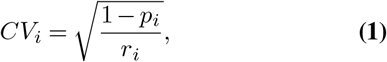

is calculated with the estimated hyperparameters. In the last step, StoatyDive normalizes the CV score by the max and min of all scores,

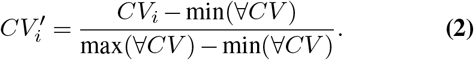

At the end, our defined CV score will range from *CV*_*i*_ = [0, ∞] and the normalized score from 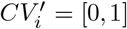, with a 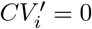 for a more unspecific binding and 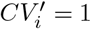 for a more specific one.

### Classification of Peak Profile

StoatyDive classifies the peak profiles in an unsupervised manner using uMAP (8) and k-means clustering (9). Yet before clustering, StoatyDive processes the peak profiles. First, the profiles are normalized based on the individual maximum and minimum read count, since we are only interested in the shape of the profiles and not in the absolute read counts (Figure 2 step 4). So assuming each peak *X*_*i*_ has *x*_1_, *x*_2_, *x*_*j*_…, *x*_*n*_ nucleotides, we normalized the peaks by 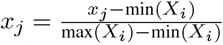. Second, the peak profiles are smoothed (Figure 2 step 4) with a spline regression (10). The step reduces the noise for each profile and distributes the data more uniformly on the current manifold. The latter is important since it is the data assumption of uMAP. StoatyDive further adds curve specific features to the processed peak profiles including: the number of maximal values, the area under the curve, and the arc length. StoatyDive applies uMAP to the final data with 5, 000 epochs, 2 components (dim = 2), a minimum distance of 0.01 and a size of the local neighborhood of 5. The dimensional reduction was optimized with some test data comprising four different sets of distributions: a uniform distribution, a linear distribution, an unimodal Gaussian distribution, and a bimodal Gaussian distribution. Subsequently, StoatyDive applies k-means clustering to the new data with 100 initializations, and maximal 10, 000 iterations. The number of clusters *k* is found by convergence of the total within-cluster sum of squares and checked with the Akaike information criterion (AIC) (11). We also tested other dimensional reduction methods such as principal component analysis (PCA), t-Distributed Stochastic Neighbor Embedding (t-SNE), and a self-organizing map (SOM). However, none of them came close to the results of uMAP (Supplementary Information: Figures 6, 7, and 8).

### Output of StoatyDive

For the peak evaluation, StoatyDive generates a plot of the CV (Equation 1) and normalized CV (Equation 2) distribution (Figure 2). The user receives a first impression of the binding specificity of the protein of interest from the CV distribution. An unspecific binder has a CV distribution close to zero. A more specific binder has a CV distribution equal to or higher than one. The CV distribution can also be used as a quality control to compare control and signal experiments. A quality breach might have occurred if the distributions of the control and signal experiment almost look identical. A control experiment should normally have a CV distribution close to zero, with only a very few binding sites showing higher CVs.

The normalized CV distribution helps to evaluate the peaks based on the individual experiments. An empirical threshold is set at a normalized CV of 0.5 (Equation 2). Binding sites with a CV < 0.5 are more unspecific than binding sites that have a normalized CV >= 0.05. The user can change the threshold. Keep in mind, the threshold for the normalized CV is relative in accordance to the individual experiment. For the peak classification, StoatyDive generates a plot of the k-means optimization and a plot of the dimensional reduction with uMAP, which can be used to readjust the number of *k* clusters if this is actually needed. The user also receives a set of example peak profiles and smoothed peak profiles of each cluster, which can be used to investigate the identified shapes. For a general trend, StoatyDive delivers average profiles for each cluster.

The final output of StoatyDive is a CVs orted table of the whole peak set, from the highest to the lowest CV. Each peak is labeled with 0, for more specific binding sites, and 1, for more unspecific sites. The table also lists for each peak the cluster number (group number) of the peak profile shape.

### Important Options of StoatyDive

The peak correction (Figure 2 step 1) can be turned off. The user can also change the translocation scheme of the peak profiles to shift them based on the maximal value (summit). The maximum translocation scheme is useful for nucleotide specific events such as truncation events in the case of eCLIP data (5). StoatyDive has also the option for a different CV score that penalizes peaks within broad plateaus. StoatyDive then adjusts the CV score of peaks that are covering a small appendage of a read stack. Furthermore, the user can provide a maximal score to StoatyDive to normalize the CV distribution (Equation 2). This option helps to compare the CV distribution between experiments in accordance to their disparate peak sizes and total amount of reads. StoatyDive also has a threshold for the normalized CV score to divide the peaks into more specific and more unspecific binding sites, which the user can change.

StoatyDive has two major parameters for the peak profile classification (Figure 2 step 6). First, the user can adjust the maximal amount of potential peak clusters identified by the k-means clustering. Yet, the final number of peak clusters will be optimized by StoatyDive. The parameter is an upper bound. However, the user has the option to force StoatyDive to use *k* specific clusters. The smoothing (Figure 2 step 4) of the peak profiles can also be adjusted by the user. The default was optimized with different test sets. Increasing the parameter (> default) might underfit the smoothing and thus lead to fewer peak clusters. A lower value (< default) might overfit and so lead to more clusters. The smoothing can also be turned off, but it is recommended to turn it on.

## Results and Discussion

### Data Preparation of SLBP and Analysis

We used eCLIP data of the histone stem-loop-binding protein (SLBP; ENCSR483NOP; GSE91802) (5). The data comprised two CLIP replicates and one size-matched input control from immortalised myelogenous leukemia cells (K562). We processed the data with the snakemake pipeline SalamiSnake (https://github.com/BackofenLab/SalamiSnake, v0.0.1) for eCLIP data. SLBP has been reported to be cytoplasmic and be present in the nucleus (6). Thus, we mapped the reads against hg38 genome with STAR (12), but also taking the transcriptome into account. We predicted potential binding sites of SLBP with PureCLIP (13), which we ran for each CLIP replicate separately, taking the input control into account. We extended the predicted binding regions by 20 nucleotides left and right because PureCLIP often underestimates the binding region. We further fused the predicted peaks from each CLIP replicate with bedtools (7) to get a robust set of predicted binding sites. We executed StoatyDive with the length normalization, the penalty for broader plateaus, and the peak profile smoothing. The complete call was: *StoatyDive.py -a peaks.bed -b reads.bam -c hg38.chrom.sizes.txt --peak_correction --scale_max 10 --border_penalty --sm*.

### CV Results Reveal Potential Quality Breach in SLBP Data

Both the CV distribution of the input control and replicate one of the SLBP data contain a lot of regions with a CV close to zero (Figure 3). In contrast, the CV distribution of replicate two is different since it had more peaks with a higher CV and thus more specific bindings such as peak 529 (Figure 1a, *CV* ≈ 5.3). Yet, some potential binding sites were more unspecific with a CV closer to zero such as peak 805 (Figure 1b, *CV* ≈ 0.006).

**Fig. 3.**
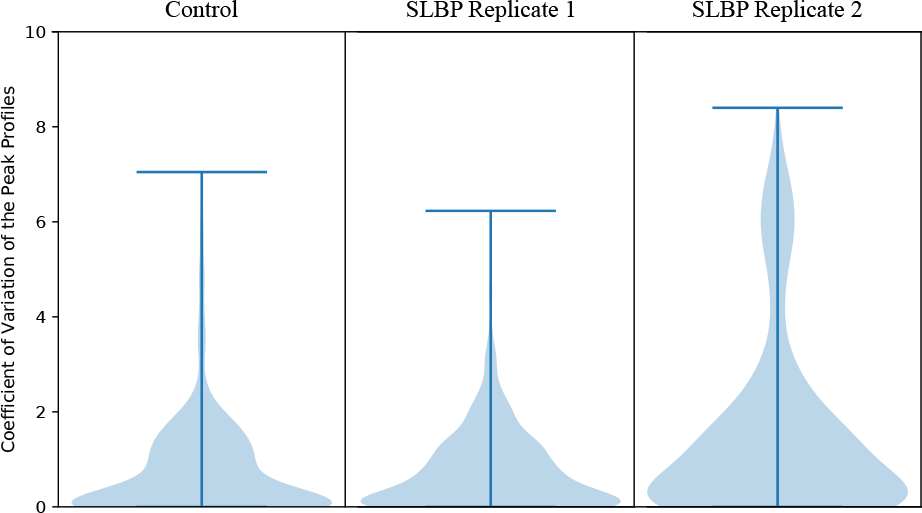
Evaluating the peak profile shapes is a quality control. (a) The distribution of the coefficient of variation of the peak profiles of the input control and replicate one of the SLBP CLIP-Seq experiment are very different to the distribution of replicate two. Thus, a quality breach might have occurred in replicate one. The user can subsequently inspect with StoatyDive more specific, sharp profiles (Figure 1a) and more unspecific, broader profiles (Figure 1b). Perhaps only one of the replicates is informative for specific patterns.

The CV distribution of the input control was expected, because ideally the control experiment contains no real binding events. But, the CV distribution of replicate one did not match the assumptions. The distribution is an indicator for a possible quality breach in replicate one. This is also striking when we compared the distribution of replicate one and two. For a downstream analysis, for example the prediction of sequence motifs, it is worth to either exclude replicate one or to investigate, why the CV distribution of replicate one was very different to replicate two. Thus given the inspection of StoatyDive, the user can now decide to investigate the unspecific binding sites of replicate one and compare them with the input control or replicate two. This helps to assess if SLBP might have protein domains that bind to RNA in an unspecific manner. The user can also test if the unspecific peak 805 might be a false positive of PureCLIP.

### Seven Different Peak Shapes in the SLBP Data

For a more detailed analysis, we classified t he p eaks o f replicate one and two with the help of StoatyDive (Figure 4). StoatyDive finds seven distinguishable peak profiles for replicate one and two. Because we assumed more unspecific sites in replicate one, we looked deeper into the profiles of replicate two (Figure 4b). Cluster two and five were clearly disparate. Groups one, three, four and six were much closer together than group two and five. Especially cluster four and six were similar. All four groups had profiles with mountain-like shapes. In contrast, cluster two and five are plateau shaped profiles. The mountains in the profiles became more broader and fuzzier from cluster three to one and then to six and four. The average, centered profiles (Supplementary Information: Figure 5 b1–7) reflected the trend of the ensemble of the individual profiles in each cluster (Figure 4 b1–7). To return to our initial examples (Figure 1), peak profile 529 was classified by StoatyDive as a small, centered mountain (Figure 4 b3), whereas peak profile 805 was a very broad profile (Figure 4 b4).

**Fig. 4.**
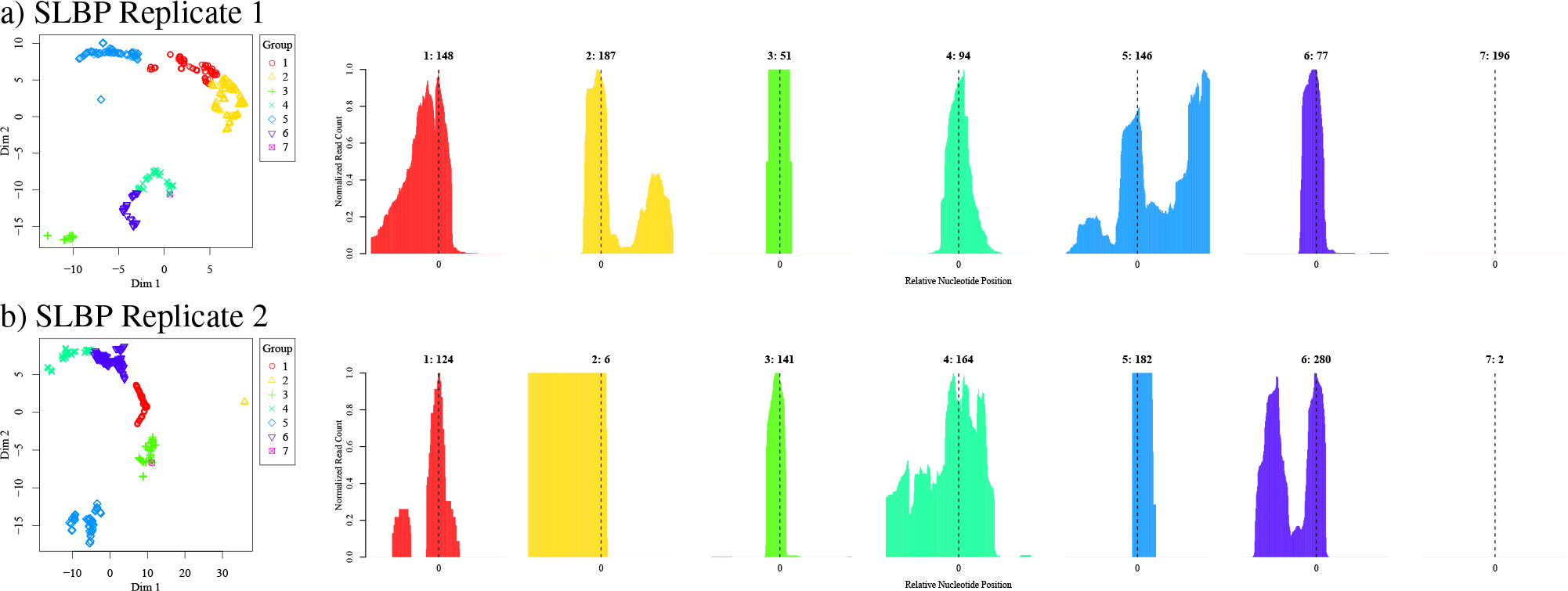
We applied StoatyDive to the SLBP data (ENCSR483NOP; GSE91802) (5). StoatyDive found seven different peak profile shapes in the data of replicate one a1–7 and two b1–7 of SLBP. One example profile of each cluster is shown with the number of peaks in each cluster in the title. The first plot shows the peak profiles in the lower dimension, when we applied uMAP. For replicate one, shape one is a broad peak profile. Shape two is a thin, spiky mountain, but with additional peaks surrounding it. Shape three is a flat plateau. Shape four is a mountain broader than shape six but thinner than shape one. Shape five is a very broad profile (cordillera). Shape six is a very thin and specific mountain. Shape seven is a constant profile. For replicate two, shape one is a small steep mountain with some small peaks surrounding it. Shape two is a very broad plateau. Shape three is a very thin and specific mountain such as peak 529 (Figure 1a). Shape four is a very broad profile (cordillera) such as peak 805 (Figure 1b). Shape five is a clear, unvarying plateau. Shape six is a broad profile with two or more clear peaks in close proximity. Shape seven is a constant profile.

It is to mention that the number of clusters depend on the optimization of StoatyDive, or the user defined value. Other proteins, experimental conditions, and methods might have different peak profile groups. Even in our example it is worth to investigate if cluster four and six of replicate two can be separated more distinctly. At this point, one could run StoatyDive again, but only with the peaks of cluster four and six.

The high distance between cluster two and five might result from the difference between the profile borders (compare Figure 4 b2 and b5). Where cluster two had profiles with lots of values in the left or right side of the peak profile, cluster five occupies the center of the peak profile.

A broader and fuzzier profile might not necessarily mean that it is an unspecific site. Perhaps some of them are just a collection of several specific peak profiles that are merged together. This could have happened because of the peakcaller model or the peak correction and extension of StoatyDive. It is worth to take these profiles and reduce the extension, and in addition run a peak deconvolution.

It is important to note that peaks shaped like plateaus might be false positives. These peaks most likely correspond to PCR duplicates that are not real binding sites. We have removed PCR duplicates during the preprocessing of the read library, but some duplicates might be still in the data. Sequencing errors in the unique molecular identifiers (UMI) are a common reason.

SLBP has been reported as an mRNA export and translation factor (6). Thus, it is worth to investigate if peaks like 529 are more informative for a translation factor than peaks like 805. That is to say, 529 might be more suited for sequence and structure predictions than peak 805. Therefore, a deeper inspection of group one, three, four and six seems promising.

### Information from Peak Profile Shapes

We made the assumption that replicate one might have more unspecific and less distinguishable profiles than replicate two based on the different CV distributions (Figure 3). Thus, we counted the number of peaks in each cluster for replicate one and two (Table 1). From 899 peaks, in replicate one we had 171 peaks being a sharp mountain shape (Figure 4 a4 and a6), 481 being a broader mountain (Figure 4 a1, a2 and a5), 51 peaks with plateaus (Figure 4 a3), and 196 constant shaped peaks (Figure 4 a7). Replicate two, on the other hand, had 265 sharp mountain shaped peak profiles (see Figure 4 b1 and b3), so 94 more than replicate one. This corroborates the assumption that replicate one has more broader and unspecific sites. Thus, replicate two had only 444 broad peak profiles (Figure 4 b4 and b6), and only 2 constant peak profiles (Figure 4 b7). Yet, replicate two had 188 peaks with plateaus (Figure 4 b2 and b5).

**Table 1.**
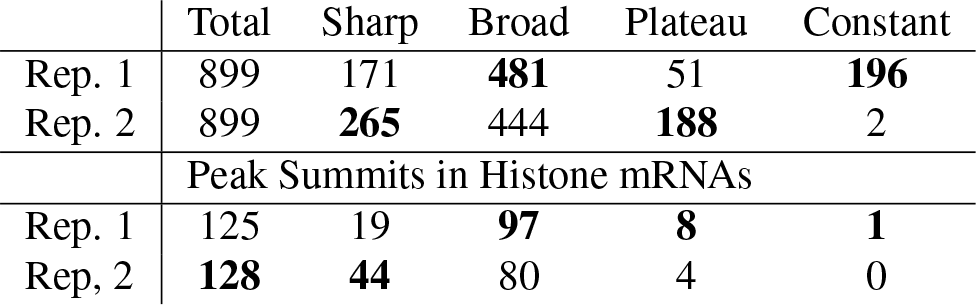
Number of peaks of SLBP for different shape groups.

|  | Total | Sharp | Broad | Plateau | Constant |
| --- | --- | --- | --- | --- | --- |
| Rep. 1 | 899 | 171 | **481** | 51 | **196** |
| Rep. 2 | 899 | **265** | 444 | **188** | 2 |
|  | Peak Summits in Histone mRNAs |  |  |  |  |
| Rep. 1 | 125 | 19 | **97** | **8** | **1** |
| Rep. 2 | **128** | **44** | 80 | 4 | 0 |

We further investigated the biological function of different peak profiles of replicate two. Since SLBP targets histone mRNA (6), we intersected known annotated mRNAs of histones (hg38, Ensembl) with the summit of the different peak profiles (Table 1). From the 899 peaks, only ≈ 14% (128 peaks) of replicate two overlapped with mRNAs of histones. Yet, of these 128 peaks almost all come from group one, three, four, and six. These groups are either spiky, or broader mountain shaped peak profiles. Only 4 peaks intersected with histone mRNAs that had a profile shaped like a plateau (Figure 4 b5). This endorses the assumption that peak profiles shaped like plateaus are mainly PCR duplicates that are less informative. The observation also suggests that broader profiles are still informative, because some of them overlapped with histone mRNAs. At this point, it is worth to investigate for downstream peak analysis tasks, such as binding sequence motif detection, how much noise each individual mountain shaped profile encompasses.

## Conclusion

StoatyDive is a powerful tool that can evaluate and classify peak profiles. It can be used in any sequencing data analysis that involves the prediction of binding sites such as CLIP-Seq, or ChIP-Seq. Within this work, we provided an example for SLBP to show the usability of StoatyDive. First, it is possible to assess the quality of an experiment such as CLIP. Second, StoatyDive assists to evaluate the binding specificity of the protein. The user should check for the normalized CV distribution. A protein that binds very specific will have a distribution concentrated around a normalized CV of one. A protein with a lot of unspecific bindings will have a normalized CV distribution around a value of zero. Third, StoatyDive helps to filter for specific and unspecific binding sites to investigate if the protein has multiple protein domains that have different binding mechanisms. A finer distinction can be made with the classification mode of StoatyDive. This helps to identify peak profiles with a specific shape and filter them based on the corresponding biological question and function of the protein. For example, a transcription factor might have more specific bindings (more spiky mountains), than a protein complex or a helicase (more broader mountains). Fourth, the results of StoatyDive can be used to validate a peakcaller (e.g., PureCLIP), that is to say, one can assess how many false positives are in the peak sets based on the shape. Different peakcaller might result in disparate peak sets and consequently different peak profile shapes.

StoatyDive is a very powerful, well documented, and easy to apply tool that refines the binding site detection in the data analysis such as CLIP-Seq. Nevertheless, StoatyDive is a very general tool. It can be used with different types of peak calling outputs and peak data types of sequencing data (e.g., ChIP-Seq, ATAC-Seq, Ribo-Seq, and others). It serves as a quality control and filtering step to select specific binding profiles, which therefore allows to improve other binding site prediction tools such as DeepBind (14), or any other subsequent analysis tasks, to increase the accuracy for the prediction.

## Acknowledgements

We are grateful to Gokcen Eraslan for his support.

## Supplementary Note 1: Average Peak Profiles

**Fig. 5.**
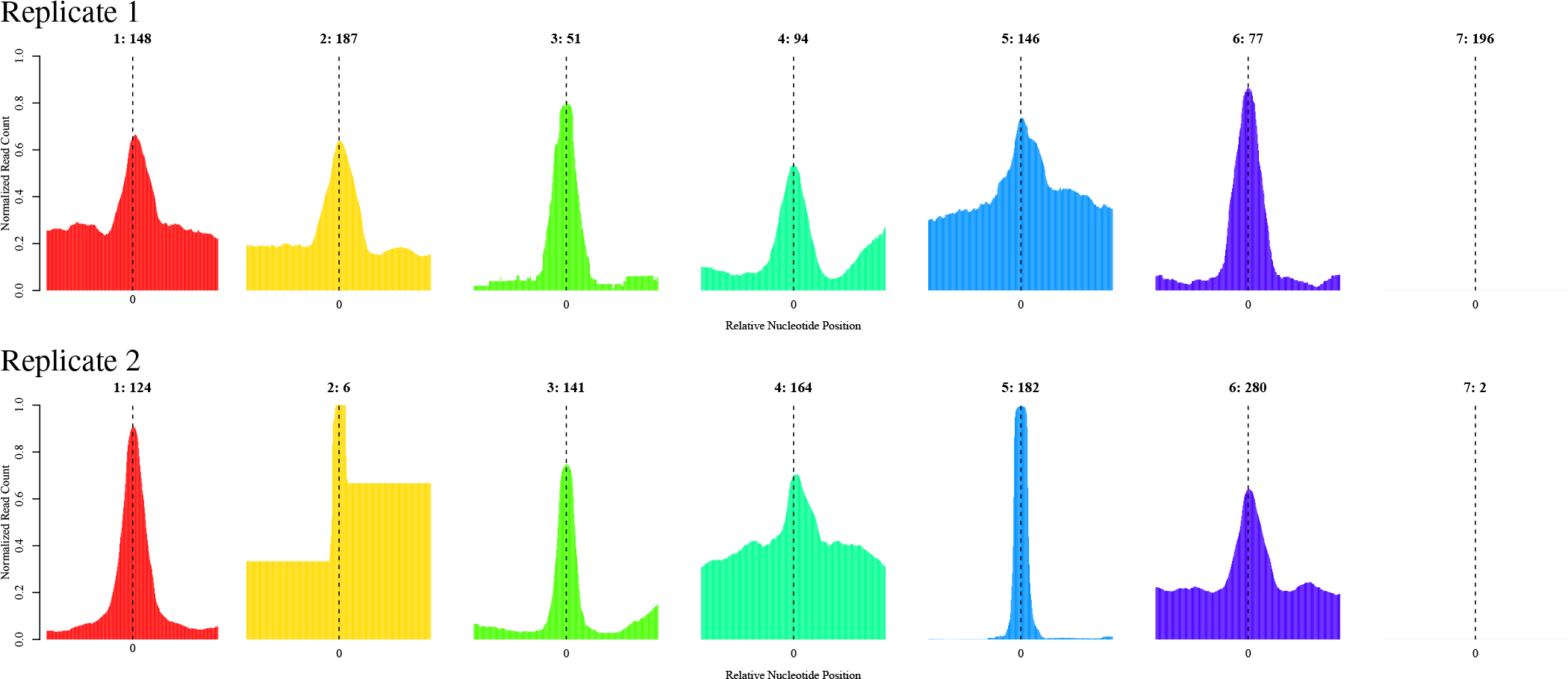
The average, centered profiles of the SLBP data of replicate one and two reflects the trend of the ensemble of the individual profiles.

## Supplementary Note 2: PCA on SLBP Data

**Fig. 6.**
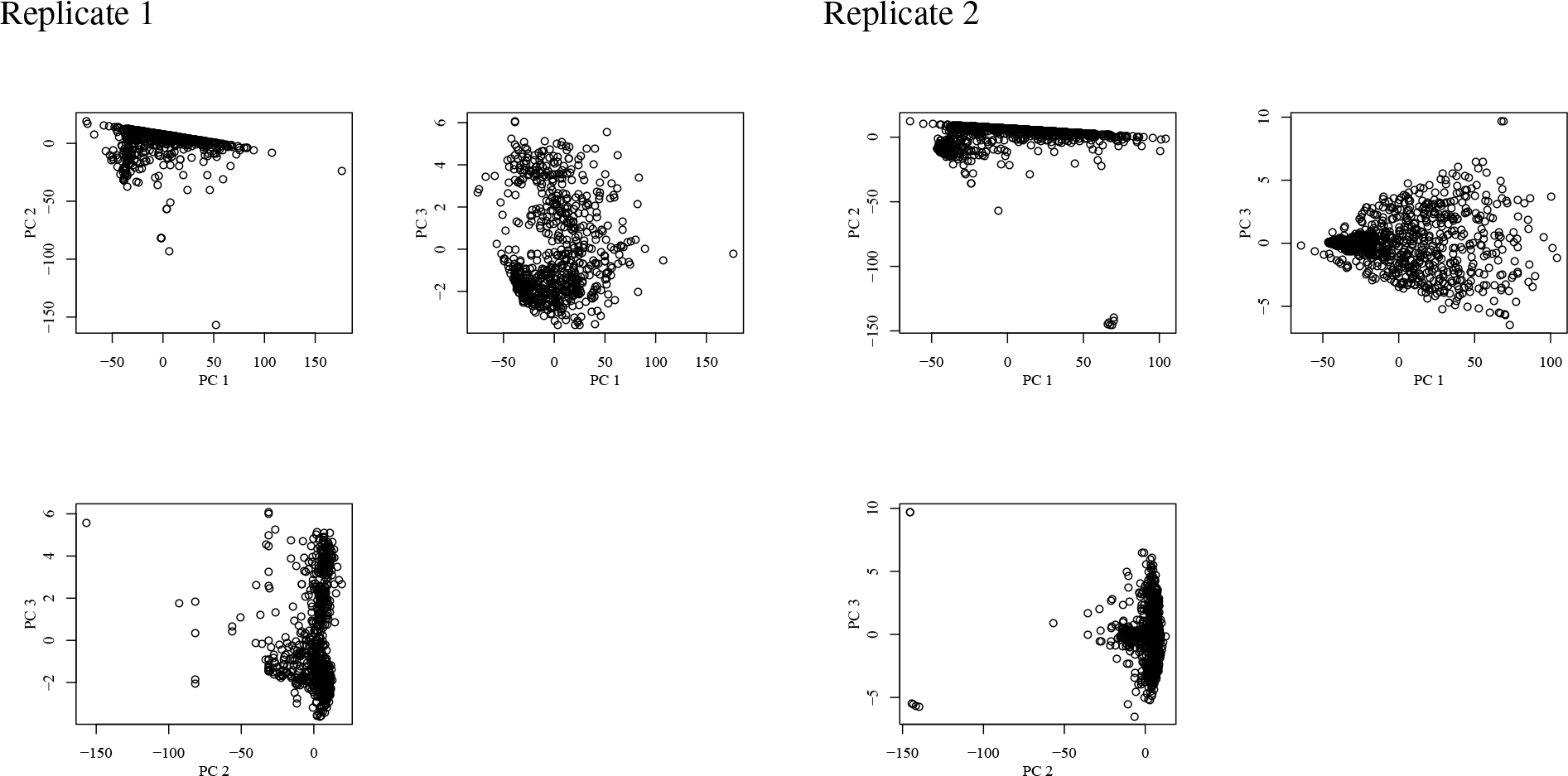
Principal component analysis (PCA) on the data that is also used for uMAP. There are no clear clusters visible after the dimensional reduction for both replicates. uMAP can clearly separate the data into more defined clusters.

## Supplementary Note 3: t-SNE on SLBP Data

**Fig. 7.**
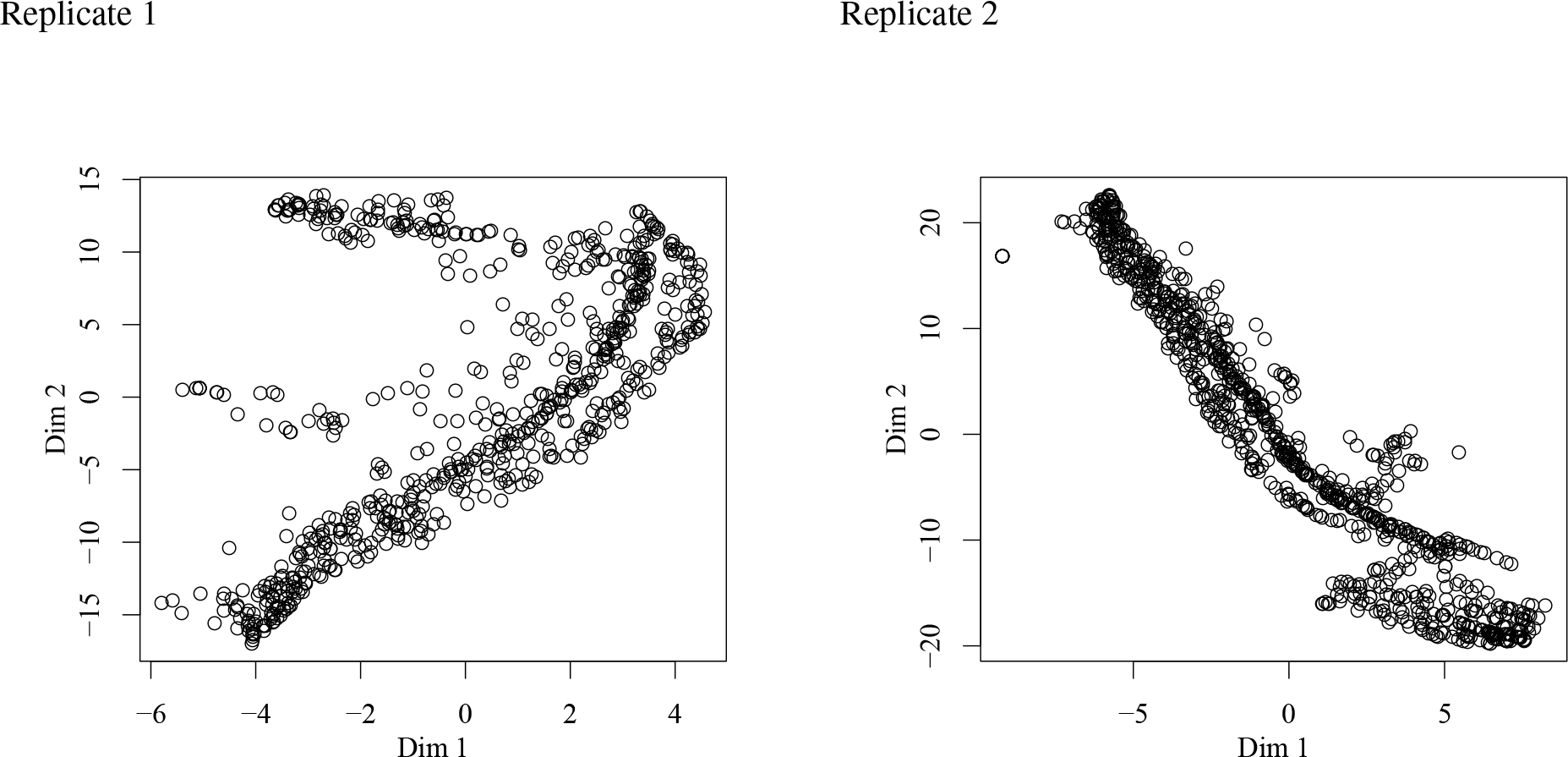
t-Distributed Stochastic Neighbor Embedding (t-SNE) on the data that is also used for uMAP. There are no clear clusters visible after the dimensional reduction for both replicates. uMAP can clearly separate the data into more defined clusters.

## Supplementary Note 4: SOM on SLBP Data

**Fig. 8.**
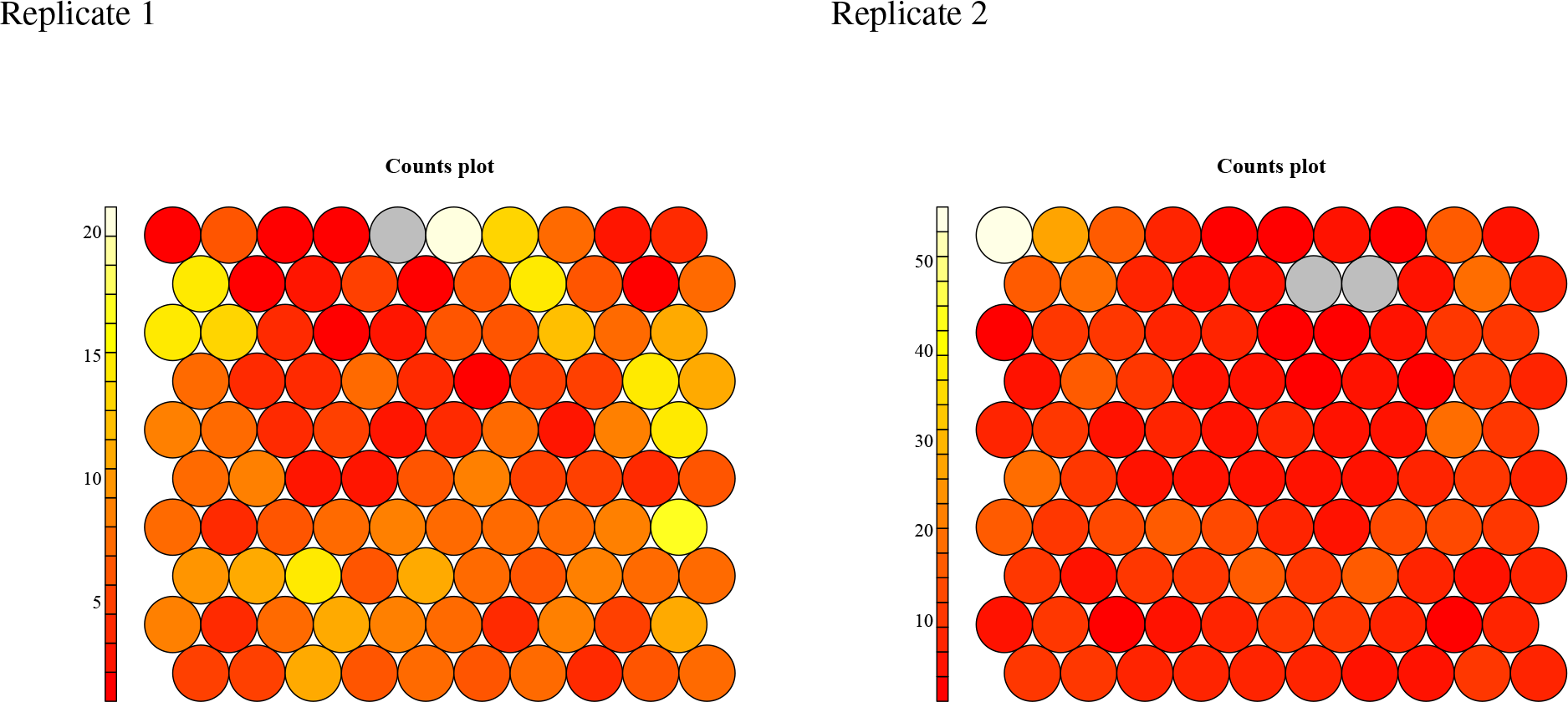
Using an optimized self-organizing map (SOM) delivers a feature layer with some high activated hidden units for replicate one and only one very high activated hidden unit for replicate two. It is hard to see any distinct clusters from the counts (activation) of each hidden unit. uMAP can clearly separate the data into more defined clusters. Furthermore it is much easier to interpret the results of uMAP, whereas an artificial neural network, such as a SOM, generates a feature layer (hidden layer) that is hard to grasp.

